# DynaMAP: Dynamic Microbiome Abundance Profiling through high density optical mapping

**DOI:** 10.1101/2024.12.05.626436

**Authors:** Sergey Abakumov, Elizabete Ruppeka-Rupeika, Xiong Chen, Mattias Engelbrecht, Inge Mestdagh, Thomas D’huys, Agata Mlodzinska, Volker Leen, Agata Szymanek, Aditya Badola, Tomas Lukes, Thomas Demuyser, Johan Hofkens, Peter Dedecker, Arno Bouwens

**Affiliations:** Department of Chemistry, Laboratory for Nanobiology, Celestijnenlaan 200G, 3000 Leuven, Belgium; Department of Chemistry, Laboratory for Molecular Imaging and Photonics, Celestijnenlaan 200F, 3000 Leuven, Belgium; Perseus Biomics, Bio Incubator 5, Gaston Geenslaan 3, 3001 Leuven, Belgium; AIMS lab, Center for Neurosciences, Faculty of Medicine and Pharmacy, Vrije Universiteit Brussel (VUB), Belgium; Department of Microbiology, Universitair Ziekenhuis Antwerpen (UZA), Edegem, Belgium; Department of Bioscience Engineering, Research Group Environmental Ecology and Applied Microbiology, University of Antwerp, Antwerp, Belgium; Bioidea, 02-991 Warsaw, Poland

## Abstract

Microbial abundance profiling is rapidly becoming an essential method in biomedical research, though it is often costly and time intensive. We present DynaMAP, a rapid approach for microbiome profiling that uses single-molecule imaging to develop an optical map of metagenomic DNA. DynaMAP achieves strain-level taxonomic profiling without requiring base-by-base sequencing and with a one-day turnaround time. In doing so, it delivers microbiome profiling that is comparable to shotgun sequencing but operates more efficiently, strongly expanding the possibility for microbiome analysis.

---

The microbiome plays a crucial and symbiotic role in maintaining the health of all plant and animal species in which it has been studied, including humans [1]. Imbalances in microbial composition, known as dysbiosis, are linked to various health conditions ranging from gastrointestinal disorders [2] to neuropsychiatric diseases [3], [4]. The presence of certain species in the human gut such as *Fusobacterium nucleatum* is furthermore associated with an increased risk of diseases such as colorectal cancer [5]. Understanding the composition and dynamics of the microbiome in normal development and in response to external stimuli is therefore essential to identify novel diagnostic biomarkers, therapeutic targets [6], and tailored interventions. Doing so with same-day turnaround times and in a cost-efficient manner would be essential to enable its application across a wide range of settings, including situations that need fast follow-up such as the detection of antimicrobial resistance in patients or fast strain-resolved microbiome profiling in the prevention of necrotizing enterocolitis in preterm infants [7].

A key challenge is the complexity and variability of the microbiome, which make the data acquisition and analysis challenging. Sequencing-based methods such as shotgun metagenomic sequencing (SMS) have advanced metagenomics and revealed host-microbe interactions, but are complex, labor-intensive, and can display biases arising from the library preparation and amplification, potentially skewing microbiome profiles from their true composition [8],[9]. The high cost of the analysis and turnaround times on the order of days or weeks also complicate its use as a widespread clinical diagnostic tool [10], [11]. This is further exacerbated by the increased sequencing depth required for the strain-level profiling that can detect low-abundance but potentially clinically important species [12]. Accordingly, methods based on base-by-base DNA sequencing struggle to deliver metagenomic data at the density and speed required to accurately understand the microbiome composition and the relations that govern it.

An alternative approach is optical mapping (OM). In this method, DNA fragments are site-specifically labeled and the spatial distribution of these labels within the fragment is visualized. When combined with optical imaging, many DNA fragments can be quickly and efficiently recorded, resulting in a procedure that is highly amenable to automation and rapid data acquisition. This enables optical mapping to achieve same-day turnaround times [13] while not requiring library preparation, PCR amplification, sequencing, and read assembly. To date, the method has been applied to the detection of structural variants within genomes of eukaryotic organisms via the sparse tagging of ultra-high molecular weight DNA at 6 or 7-mer recognition sites [14], for bacterial strain identification using fragments over 100 kb in size [15], to discriminate between a limited number of strains in a sample [16], and to detect antimicrobial resistance [17] [18]. However, the current requirement for long DNA fragments, the low labeling densities, and/or the small set of strains that can be included in the analysis limit the use of common samples such as stool due to their high nuclease burden [19], [20] and high species variety.

The fast performance of optical mapping marks it as a promising tool for metagenomic profiling, provided that it can achieve the required information density to differentiate between complex microbiomes using labeled DNA fragments of limited length. Here, we introduce DynaMAP, a method that uses high-density fluorescence DNA labeling combined with fast imaging and powerful data analysis in order to assess the presence of DNA from thousands of distinct species within a single sample. We show that this method can achieve comparable performance to whole-genome sequencing but achieves a faster turnaround time and lower cost.

Figure 1a provides a detailed overview of the DynaMAP process. The measurement begins with a high molecular weight (HMW) DNA extraction, optimized for stool samples, that produces fragments with a median length of 40 kbp (Supplementary Figure S1). These are then labeled by introducing fluorophores at TCGA sites using the methyltransferase enzyme *M*. *TaqI*, such that there is on average one label every 256 DNA bases. These fragments are then combed onto a flat surface and imaged using single-molecule fluorescence microscopy, resulting in a distinct optical map for each fragment. A pattern-matching algorithm then aligns each optical map with a large variety of reference genomes and determines the originating species. Processing each sample takes approximately 6.5 hours, including 2 hours for DNA extraction and labelling, 1 hour for imaging, and 3.5 hours of processing time to analyze a total of around 6 gigabases of DNA (around 140000 optical maps) using a reference database containing 4992 distinct genomes.

**Figure 1:**
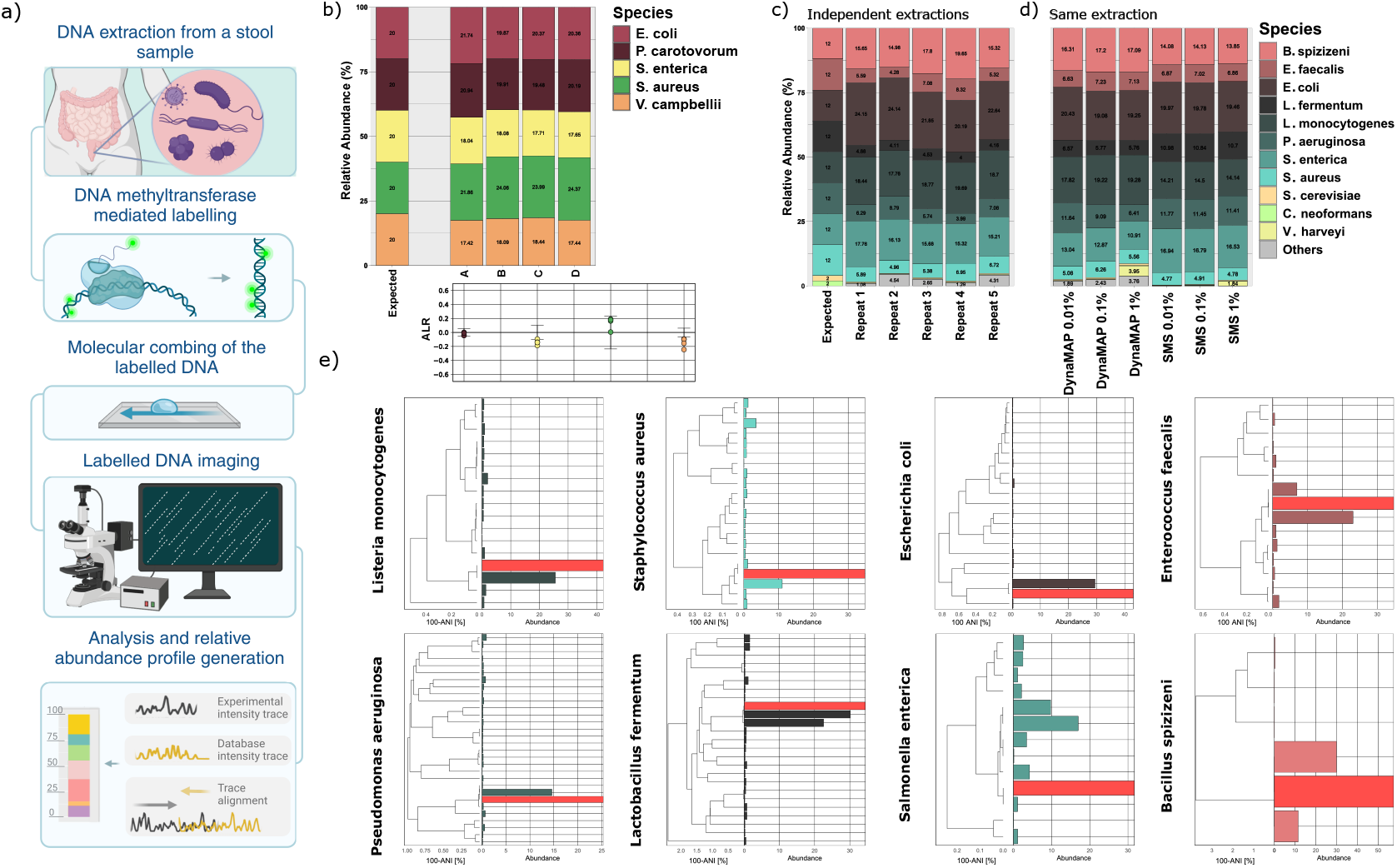
(a) Overview of the DynaMAP workflow, including DNA extraction, methyltransferase labeling, molecular combing, imaging, and profile analysis. (b) Performance on a mock DNA mix: each relative abundance plot represents an independent labeling of the same mix. The inset represents ALR-transform with respect to *E. coli*, with error bars indicating possible variation of ALR as predicted from concentration measurements. (c) DynaMAP relative abundances from five technical repeats of D6300 extraction, showing repeatability and alignment with reference values. (d) Limit of detection testing with spiked *V. harveyi* at 1, 0.1, and 0.01%, suggesting a lower bound of 0.03% for DynaMAP’s FP rate. (e) Strain-level abundance estimates for ZymoBIOMICS D6300 strains, showing taxonomic proximity and relative abundance (MLE).

We evaluated DynaMAP’s performance by applying it to a mixture containing equal amounts of purified DNA obtained from five separately cultured bacterial species. To minimize the effects of lysis bias, DNA fragmentation, and reference genome quality, we selected species that offer comparatively straightforward DNA extraction and high-quality genome information. The DNA was extracted using a specialized kit designed for the extraction of high-molecular weight DNA from pure cultures of bacterial species. We then subjected mixtures of equal amounts of DNA to the full DynaMAP procedure in four independent replicates (A–D in Figure 1b), analyzing the results with a reference database containing only the genomes of those five species. The results showed consistent species recovery, with relative abundances between 17– 23% for each species. We also looked at the absolute log-ratio transform (ALR) of each species relative to *E. coli* and conducted a one-sample t-test (p = 0.05) to determine if certain species were over- or underestimated compared to this bacterium (see Materials and Methods, Sample preparation). This analysis found no significant deviation for two of the four species, though did reveal significant differences for *V. harveyi* and *S. enterica* for which the overall abundance was estimated as 18.2% and 17.95% rather than the expected 20%. We repeated the analysis using the full set of 4,992 reference genomes, finding that the average cumulative abundance of the fragments across all repeats attributed to species not present in the sample was 3.67±1.51%. (Supplementary figure S3). We defined the false positive abundance (FPA) as the average detected abundance of those species that are not present in the sample. This value was consistently very low at 0.034±0.007%. Overall, this experiment suggested that DynaMAP could successfully identify distinct bacterial species with good accuracy.

We then applied our approach to the ZymoBIOMICS microbial community standard D6300, a mixture of three Gram-negative bacteria, five lysis-resistant Gram-positive bacteria, and two lysis-resistant yeasts. This standard is commonly used to assess metagenomic workflows, including the effect of lysis bias. We compared the expected ground-truth composition, as provided by the manufacturer, with five repeat experiments of the full DynaMAP pipeline, which included independent lysis, labelling, imaging, and analysis with the full 4,992 species reference database, performed on separate D6300 aliquots. The results, shown in Figure 1c, compare the measured and expected abundances, aggregated at the species taxonomic level, for 5 repeated extractions. The measured abundances of the 8 bacterial species align with the expected values, with no apparent gram-stain bias in the extraction process, which is a common concern in metagenomic studies [21]. All species were successfully detected, though four species, *Pseudomonas aeruginosa, Lactobacillus fermentum, Enterococcus faecalis*, and *Staphylococcus aureus*, were underrepresented. The low abundant yeasts, *Saccharomyces cerevisiae* and *Cryptococcus neoformans*, present in the mock were also underrepresented, which may be due to lack of enzymes tailored to yeast lysis. The underrepresentation of *S. aureus* and *E. faecalis* may arise their resistance to lysozyme [22]. The lysis bias introduced by our HMW extraction requirement is comparable to extraction biases of various commercial kits as observed in [23], and produces repeatable profiles as can be deduced from Figure 1c.

The analysis results also contained the presence of species that were not present in the sample, though the average false abundance was likewise very low, ranging from 0.013±0.005% to 0.038±0.016%. The average total abundance of fragments across all repeats that were matched to incorrect reference genomes was 2.77±1.45%. Based on these findings, we could estimate the limit of detection (LOD) for optical mapping to be in the range of 0.028% to 0.086%, using the definition of LOD as being 3σ above the average background level. To verify this in practice, we spiked the D6300 standard with low concentrations of *Vibrio harveyi* strain BB120 DNA (0.01, 0.1 and 1%), which originally was absent from the D6300 mixture, and obtained total measured abundances were 0.027%, 0.23%, 3.95% (Figure 1d), where 0.027% is below the level of detection as found for Figure 1c. In addition, the extracted DNA from each spike-in was subject to a separate evaluation by SMS. The spike-in abundances measured by SMS were 0.016%, 0.17%, 1.84% respectively. Our observed LOD is somewhat higher than the detection limit of SMS (∼ 0.01%) within this experiment, a value that has also been reported in literature [24]. Overall, our DynaMAP approach achieves similar results as SMS-based analysis, though with a somewhat higher limit of detection. Furthermore, the comparison of microbial compositions with the results from SMS showed an excellent correspondence between two methods, yielding errors in the relative abundances of about 1-5% between both methods (Figure 1d). Based on these considerations, we applied a 0.1% abundance cutoff on the DynaMAP analysis, meaning that we do not report species or strains with abundances below this level determined by DynaMAP to eliminate all false positive detections from complex samples (Figure 2).

**Figure 2:**
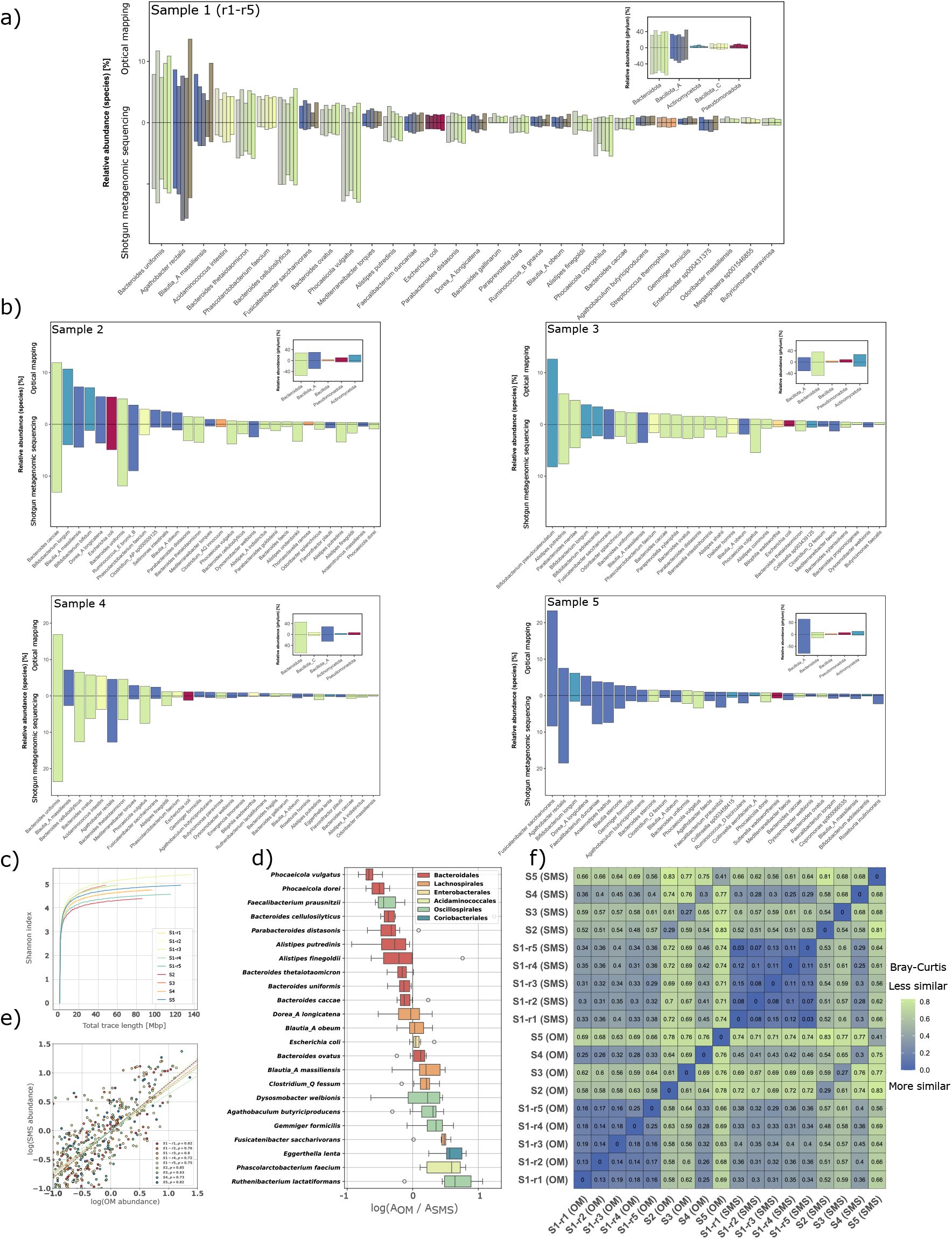
(a) Comparison of DynaMAP optical mapping and shotgun sequencing -based gut microbiota compositions from 5 repeats of a sample, with species-level data and phylum-level insets. (b) Comparison for 4 additional samples, showing high concordance between methods. (c) Rarefied Shannon index for optical mapping, saturating at 60-80 Mb classified data. (d) Log-transformed ratios of optical mapping (OM) to SMS abundances, suggesting conversion potential between methods. (e) Log-log scatter plot of relative abundances with Spearman correlations per sample. (f) Bray-Curtis distance matrix for all samples, showing repeatability across 5 technical repeats and low distance between methods for samples from the same donor.

The performance of the DynaMAP analysis suggested that it could be applied to resolve distinct strains within a sample. To verify this, we performed an alignment of the DynaMAP results obtained on the ZymoBIOMICS standard to a database containing a large number of strains of only those species that were identified in the mixture (ranging from 7 for *B. spezizenii* to 4,428 for *E. coli*; Materials and Methods). This enabled us to efficiently profile hundreds to thousands of similar strains simultaneously (see Supplementary Table S2). The results in Figure 1e illustrate that DynaMAP resolved the expected strains within the ZymoBIOMICS D6300 standard, up to a very average nucleotide identity (ANI) level of over 99.5% similarity. Additional simulations (Supplementary Note 2) show that while longer fragments (up to 120 kb) improve accuracy for strains with up to 99.8% ANI, even 40 kb fragments reliably distinguish strains at 99.5% ANI, the threshold where strains are often considered to be identical.

This performance results from DynaMAP’s use of long-range mapping, which is sensitive to structural variation. Such strain identification is more difficult using short-read sequencing, which struggles with strain-level accuracy [25]. As a result, DynaMAP is a powerful tool for detailed microbial profiling and strain-specific tracking in complex samples, supporting its application in pathogen tracking and other microbiome research that requires reliable strain differentiation.

Lastly, we examined the performance of our method on fecal microbiomes from five different commercially sourced human stool samples. One of the samples was assayed over 5 full independent repeats and also submitted for SMS analysis to gauge the reproducibility and correspondence of the determined microbiome compositions. Figure 2a shows the relative abundances for the five repeat analyses using both DynaMAP and SMS, while Figure 2b displays the DynaMAP results for the other samples at the species (main) and phylum (inset) levels. As can be seen from this data, both the DynaMAP and SMS methodologies delivered results with broadly comparable abundances and similar precisions (repeatability). We compared the similarity of each analysis replicate using Bray-Curtis distances (Figure 2f), revealing excellent precision though somewhat higher for SMS than for DynaMAP. We further performed a rarefaction experiment and calculated the alpha diversity (specifically the Shannon index) as function of subsampled number of kilobases (Figure 2c). These saturate after approximately 50 Mb, showing that this amount of classified genomic material is sufficient to retrieve the core microbiome profile.

Some species showed different abundances between the OM and SMS analysis, which we quantified using the log-ratio (Figure 2d). One notable example is *Phocaeicola vulgatus*, which was notably underestimated by OM possibly due to its high genome plasticity [26] that causes sequence divergence and hinders strain assignment. We suspect that high genome plasticity within the *Bacteroidales* order [27] similarly affects other genera, like *Bacteroides* and *Alistipes*. Because relative abundances are calculated in proportion to total detected taxa, the underrepresentation of *Bacteroidales* artificially inflates the relative abundance of *Lachnospirales* and *Oscillospirales*. However, the absolute accuracies of the methods are difficult to assess since neither SMS nor OM are artifact-free. If required, however, this type of dataset could be used to define species-specific scaling factors that allow reweighing of the OM data to match the likely expected result of SMS.

In summary, we have presented DynaMAP, a new method to analyze microbiome compositions in a fast and scalable way. This is the first demonstration of the method on samples of such high complexity. We demonstrated its repeatability, evaluated potential biases, and compared it with shotgun sequencing. Our results indicate a strong correspondence between the methods, establishing optical mapping as a faster and resource efficient alternative to shotgun sequencing for metagenomic profiling. We show that optical mapping is a rapid and effective method for strain resolution metagenomic profiling, with potentially ground-breaking implications for microbiome research, and many other applications of the technology. For example, it could make routine microbiome profiling in the clinic feasible and enable microbiome profiling part of standard health check-ups. DynaMAP can lead a path toward the rapid acceleration of the pace (and accessibility) of microbiome profiling and lower its cost dramatically, thereby democratizing this important new aspect of health and wellness monitoring.

## Materials and Methods

### High molecular weight DNA extraction

At first, a 100-400 mg stool sample was submerged in DNA/RNA shield and stored at 4°C prior to extraction. Subsequently, the sample was thoroughly homogenized for 5 min at 1500 rpm on a VM800 Multitube vortexer (JOANLAB). A 350 µl aliquot of fecal suspension was then transferred to a DNA LoBind tube and supplemented with 800 µl NaCl 0.9%. Thereafter the fecal sample was centrifuged at 650 x g for 2 min for macromolecular debris removal. The supernatant was removed and transferred to a clean DNA LoBind tube. A second centrifugation step at 7200 x g for 1 min followed. The resulting fecal cell pellet was kept, and the supernatant removed and stored at 4°C for downstream use. The pellet was dissolved in 1 ml PBS and centrifuged again at 650 x g for 2 min (second debris removal. The supernatant was kept and transferred into a clean DNA LoBind tube and again centrifuged at 16000 x g for 1 min. In the case of cell samples, the cellular suspension was centrifuged at 16000 x g for 1 min, excluding previous washes. The supernatant was discarded, and the fecal cell pellet was resuspended in 40 µl PBS (pH 7.4). To the fecal cell suspension, 100 µl STET buffer (8% sucrose, 50 mM Tris-HCl, 50 mM EDTA, 5% Triton X-100, pH 8.0, 0.2µm filtered), containing 6 mg Lysozyme chloride form from chicken egg white (Sigma-Aldrich, Merck) and 1 mg Achromopeptidase (Fujifilm Wako Pure Chemical Corporation) was added. The suspension was vortexed 10 times at maximum speed (3200 rpm) and incubated at 37°C for 30 min at 1400 rpm. Subsequently, the lysis mix was supplemented with 20 µl Proteinase K (200 units/ml, NEB), followed by 10 vortexing steps at maximum speed (3200 rpm). The microbial cell lysate was then incubated at 55°C for 10 min at 900 rpm. Next, 160 µl of 6 M guanidinium hydrochloride in TE buffer (10 mM Tris, pH 8.0, 0.1 mM EDTA) was added and the suspension subjected to incubation at 55°C for 10 min at 900 rpm. Finally, the supernatant stored upstream in the extraction workflow was added back to the lysate. Chloroform-isoamyl alcohol in a 1:1 ratio was added and the suspension centrifuged at 10000g for 1 min. The top layer or aqueous phase (300 µl) was collected, and to it, 400 µl ice cold isopropanol was added. The suspension was centrifuged at 16000 x g and 4°C for 15 minutes to precipitate the extracted microbial DNA, and supernatant was discarded. After washing the DNA pellet with 80% ethanol, samples were incubated at 50°C for 2 min to remove residual ethanol wash solution. Thereafter, 50 µl TE, low EDTA was added, followed by 30 min incubation at 50°C, until complete dissolvement of the DNA pellet. Once resuspended, an additional automated purification of the DNA was performed on a KingFisher Flex instrument (ThermoFisher) using magnetic beads. Finally, DNA is eluted from the beads at 50°C in 75 µl TE (pH 8.0, low EDTA).

### Sample preparation

To gauge the performance of optical mapping in absence of other biases, a calibration mixture of 5 species was prepared. We restricted our mixture to only 5 species as to minimize the possible error on the relative abundances that can be introduced by pipetting. The HMW DNA from 5 bacterial strains: *Pectobacterium carotovorum ATCC 15713, Salmonella enterica ATCC 14028, Escherichia coli K12 MG1655, Vibrio campbellii ATCC BAA 1116* and *Staphylococcus aureus ATCC 12600*, was extracted with the Circulomics kit (PacBio) according to the manufacturer protocol for gram-positive and gram-negative bacteria. For *S.aureus*, we’ve added lysostaphin (10 mg/ml final concentration). Subsequently, DNA concentration was measured by Qubit essay separately 5 times for each extraction according to the protocol provided by the manufacturer with the final concentration being the average of the five measurements. The variances obtained from the concentration measurements were used to calculate the variance of hypothesized distribution of ALR with respect to *E*.coli (see Supplementary Material S1). For each of the log-ratio transformed abundances, we’ve performed a t-test at p-value of 0.05 to a hypothesized distribution with aforemention variance and zero mean to determine whether our observations can be explained by variances due to the pipetting error.

The D6300 cellular mock was extracted according to the protocol in Materials and Methods: High Molecular Weight DNA extraction, using only the part of the protocol corresponding to the extraction from cellular samples.

Stool samples were purchased from Innovative Research, each containing approximately 10 g of fecal matter. Samples were stored at -50 degrees prior to extraction. For each extraction, 200-500 mg of fecal matter was dissolved in DNAShield and stored overnight at 4 degrees before extraction protocol was carried out. For samples 2-5, a single extraction was performed. For the first sample, we performed 5 separate extractions, using separate stool aliquots.

### DNA labelling, deposition and imaging

The extracted microbial genomic DNA was subjected to enzymatic labelling using *M. TaqI* methyltransferase and a synthetic, fluorescent SAM analogue [28]. A labelling mixture is composed of 100 μM cofactor, 0.175 μg/ μl enzyme, 100 ng DNA and 1x rCutSmart buffer (NEB) and incubated for 30 min at 60°C. Subsequently, 1 µl of Proteinase K (800 units/ml, NEB) was added, and the mixture was incubated at 50°C for 30 min. The labelled microbial DNA was then again purified on a KingFisher Flex instrument (ThermoFisher) as discussed before, with the only difference that elution was achieved in 175 µL elution buffer. Finally, 2-4 μl of purified DNA was diluted with distilled water to a final volume of 9 μl. Before DNA deposition, 1 μl of MES buffer (pH 5.6, 0.5M) was added to lower the pH between 6-7, and the 2.7 μl of DNA solution was deposited on Zeonex covered slide. The slides were prepared according to [29]. The droplet was dragged using a razorblade, with a height positioned around 0.8-0.85 mm above the coverslip (see Supplementary Figure S4).

The slides, with the exception of home-mode mock sample, were mounted onto a plexiglass substrate and attached to the stage of the Nikon Ti2 Eclipse microscope. On top of the sample, 200 μl of perfluorodecalin (Merck) was added. Imaging was performed with a Nikon CFI Apochromat TIRF 100XC Oil (NA 1.49), and the sample was scanned with a Nikon perfect focus autofocus system, illumination with L6cc Oxxius laser combiner at 561 nm, and an exposure time of 80 ms per tile and 30 mW output power after objective, with a tile size of 1536×1536 pixels (virtual pixel size of 78.6 nm) imaged on PCO Edge 2.0 sCMOS camera. The home-mode mock sample was scanned using a Zeiss Axioscan system, with a Zeiss Apochromat objective (1.4 NA) and a software autofocus system on a Hamamatsu ORCA Flash 4.0 camera with a tile size of 2048×2048 pixels and a virtual pixel size of 108 nm. Multiple slides were scanned per sample, depending on the deposition density of the DNA onto the substrate. The amount of scanned area per single scan was fixed to 55.8 mm^2^, so that the entire deposition surface can be scanned, and the average time per scan was 26 minutes. With a deposition density ranging from 18-43 Mb/ mm^2^, this yields a data throughput ranging from 55 Gb – 132 Gb per day, or 9 to 22 metagenomes, given 6 Gb of data per metagenome, without further multiplexing.

### Image processing

The obtained microscopy images required pre-processing to extract the information contained within the linearly deposited DNA fragments. This pre-processing step yielded a mask that can be applied onto the original image to extract the optical maps. First, all pixel values within the images were clipped at a threshold corresponding to the 0.1% of the upper limit of the histogram. Next the image was Fourier transformed, and x and y coordinates of pixels passing the threshold within the Fourier transform were used for subsequent orthogonal regression analysis. The error obtained from the fit was used as a threshold for discarding bad quality images, as these contained improperly stretched DNA or other artifacts and the angle from the regression coefficient was utilized for smoothing of the optical maps along their orientation direction. This last step was done by cropping the Fourier transform orthogonal to the direction of the optical maps, as shown in the Supplementary Note 3: Figure S6. In the following step, we’ve obtained the background present in the image by down sampling it by a factor 3, applying a median blur with a kernel size of 6 and subsequently up sampling it to the size of the original image. This background was then subtracted from the filtered image, and the image was re-scaled to the 8-bit range. Finally, we’ve binarized the resulting image at a white value of 240, yielding a binary mask that could be used for trace detection. To extract the optical maps, we first detected and labeled all connected components within the mask, discarding areas that are too thick (above 8 pixels) or too small (total area size below 90 pixels). Then, we’ve joined the areas according to our imposed empirical rules (see Supplementary Note 3: Figure S7). The final one-dimensional optical map is extracted by interpolating the intensity along the direction parallel to the longest axis of the detected area. Only optical maps exceeding the minimal size of 31 kb were used for analysis, and the size distribution of optical maps obtained from various samples can be found in the Supplementary Figure S1.

### Cross-correlation processing and reference database construction

The optical maps were first pre-processed before alignment. In the data processing phase, each of the optical maps was first locally normalized, with a window of 10 kb [30]. In order to generate the reference trace, we performed an *in-silico* digestion by TaqI, detecting all the potential labelling sites for each of the genomes. Next, we transformed the base-pair position to the distance map, by multiplying it with the 0.34 nm, the physical size of the base-pair and a stretch factor. The stretch factor follows a distribution, ranging from 1.64-1.78 (see Supplementary Figure S2), therefore, 10 references were generated for each of the stretch factors in that range. Then, we convolved the obtained sparse representation of our bacterial genome with a gaussian peak, for which the full width at half maximum corresponds to the FWHM of the experimental PSF. Lastly, similarly to our experimental traces, we locally normalized the reference traces with a window of 10 kb. Despite various possible speed-ups of the cross-correlation calculation, this was not yet fast enough for timely analysis of our data. We modified the cross-correlation calculation, using a multi-scaled approach, a graphical representation and explanation of which can be found in the Supplementary Note S4. Once the matches are found in the database, the relative abundances are calculated as the sum of the lengths of all matches (in kb) corresponding to a certain genome, divided by the total sum of the length of all matched optical maps.

A custom database had to be constructed to match the measured traces to reference genomes. The database included bacteria and archaea present in the human gut, including almost 5 thousand genomes of both common and rare occurrence. The source of the database is the NCBI database [31] and RefSeq and Genbank genomes, and the UHGG v2.0 database [32], including Metagenome-assembled genomes - MAGs. Genomes in the database are annotated to the GTDB version 214 [33] taxonomy using the GTDB-Tk v2 software [34]. Genomes in the database passed several quality criteria including a minimum contig/scaffold length of 200kbp and TCGA label density per 1kbp > 1. Additionally, only genomes with completeness > 90% and contamination < 5% were included in the database, with minor exceptions if the reference/representative genome was the only one available in the NCBI database. The database includes 34 phyla, 940 genera, 2,805 species and 4,992 strains.

### Strain abundance estimation

The lack of long-range information in short-read sequencing complicates strain level abundance estimation, requiring high coverage of the target genome [25]. We reasoned that the long-range information present in the optical maps captures structural variants that could permit easier identification of the strain of origin. To estimate the strain level abundance, we’ve performed a probabilistic estimate of the abundance through maximum likelihood estimation. The majority of algorithms that are used to distinguish strains are based either on base-by-base alignments of reads, or on single nucleotide polymorphism (SNP) profiling. Since optical mapping has no access to single base pair information, we’ve opted for a probabilistic approach. In our case, we treated the vector of normalized abundances 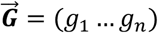 of individual strains together with matrix of trace scores ***P*** assigned to them as probabilities. From [35], it is found that by maximizing the log-likelihood,

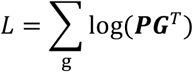

the abundance at the strain level can be inferred. To obtain the MLE, we’ve performed particle swarm optimization, subject to normalization constraint for the abundance vector ∑_*i*_ *g*_*i*_ = 1, using PySwarm with reflective boundary conditions. First, we’ve performed tests on simulated data, as shown in the supplementary (Supplementary Note S2) on strains of *Escherichia coli*. These results showed that using our approach, we are able to handle a relatively large number of strains, with ANI’s ranging from 95-99.7%. Beyond 99.8% ANI however, the true strain abundance could no longer properly be recovered and required larger fragment length for proper estimation (see Supplementary Figure S5). To verify our approach on experimental data, we’ve created strain level databases for species present in the ZymoBIOMICS D6300, by pooling all available strains from NCBI, with a completeness level of ‘complete’, and excluding any references that do not reach a 200 kb threshold in length. Next, we’ve aligned our experimental traces to this database, allowing for multiple matches, and selected only the fragments above 50 kb for strain abundance estimations. Subsequently, all the strains with MLE calculated abundance level of <0.1% were removed, and the abundances were re-normalized. To interpret results of cross matching between various remaining strains, we’ve calculated an all-to-all ANI separately for each species using FastANI [36] resulting in a similarity matrix and used the 1-ANI value as a distance matrix for hierarchical clustering. From the output of hierarchical clustering, we’ve constructed a dendrogram corresponding to ANI based taxonomy of the strains in our database. To visualize the dendrogram, we’ve only plotted the branch containing the true positive ZymoBIOMICS strain and neighboring strains for each of the species, which was visualized together with the MLE calculated abundances in Figure 1e.

### Shotgun pipeline

The sequencing was performed at Eurofins, on Illumina NovaSeq PE150 platform, obtaining ∼3 Gb of 2×150 bp reads per sample. First, the quality of raw reads and the presence of adapters from the Illumina machine were assessed using the FastQC program. In the next step, raw reads were classified against the human reference genome GRCh38.p13 (GCF_000001405.39) using the kraken2 software [37]. Reads classified to the human reference genome were removed from the further analysis, and the remaining reads were trimmed using the Trimmomatic program [38] and default parameters - the average Phred quality in a 4-nucleotide window > 30, and removal of all sequences < 30 nucleotides. Reads that passed the quality analysis were then classified using kraken2 algorithm against a previously built and indexed custom database. The database included almost 5 thousand bacterial and archaeal genomes annotated with the GTDB taxonomy version R214 [33]. The classification included a confidence threshold of 0.6. After the reads were classified, taxonomic assignment was performed at each level using the Bracken software [32] with default parameters. The identified microbiome data were used for further analysis and visualization using the R software.

## Acknowledgements

We would like to acknowledge input from Pierre Henri Ferdinand and his review of the manuscript. Furthermore, we would also like to acknowledge Rachel Kraut and her help in revision of the manuscript. Additionally S.A acknowledges funding from VLAIO Baekeland mandate HBC.2021.0848. J.H. acknowledges financial support from the FWO (Fonds voor Wetenschappelijk Onderzoek, grant number G0C1821N), the Flemish Government through long-term structural funding Methusalem (CASAS2, Meth/15/04) and the MPI as a fellow.

## Author contributions

A.Bo, V.L, J.H and P.D designed and supervised the research/experiments. S.A, E.R.R, M.E., T.D’h and I.M. performed experiments and data acquisition. X.C performed co-factor synthesis. S.A performed the data analysis of optical mapping data. A.M., A.S. and A.Ba. performed the bioinformatics analysis of shotgun data. T.L., M.B., A.Ba and S.A. provided software for optical mapping data analysis. S. A, E.R. R and A.Bo. wrote the manuscript and prepared the figures. A.Bo., P.D., J.H., T.D. and V.L. reviewed the manuscript.

