## Supplementary material for "DynaMAP: Dynamic Microbiome Abundance Profiling through high density optical mapping": DynaMAPDraftSupplementaryBioarxiv.docx

Belgium,

^2^Department of chemistry, Laboratory for Molecular Imaging and Photonics, Celestijnenlaan

200F, 3000 Leuven, Belgium,

^3^Perseus Biomics, Bio Incubator 5, Gaston Geenslaan 3, 3001 Leuven, Belgium

^4^ AIMS lab, Center for Neurosciences, Faculty of Medicine and Pharmacy, Vrije Universiteit Brussel (VUB), Belgium

^5^ Department of Microbiology, Universitair Ziekenhuis Antwerpen (UZA), Edegem, Belgium

^6^Department of Bioscience Engineering, Research Group Environmental Ecology and Applied Microbiology, University of Antwerp, Antwerp, Belgium

^7^ Bioidea, 02-991 Warsaw, Poland

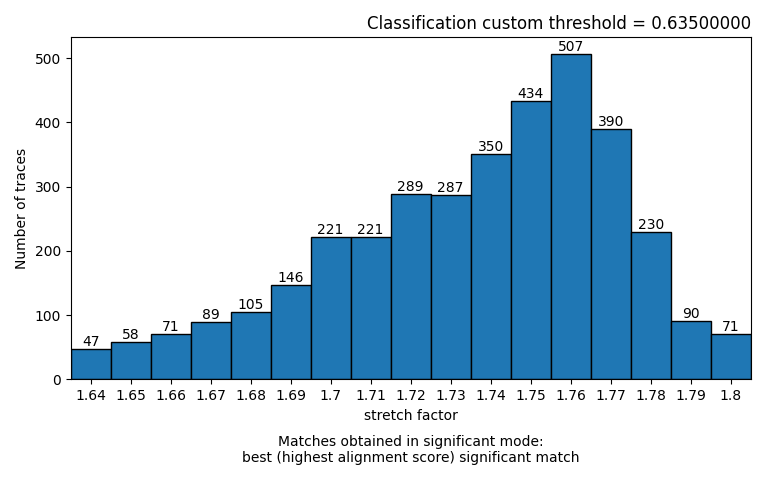

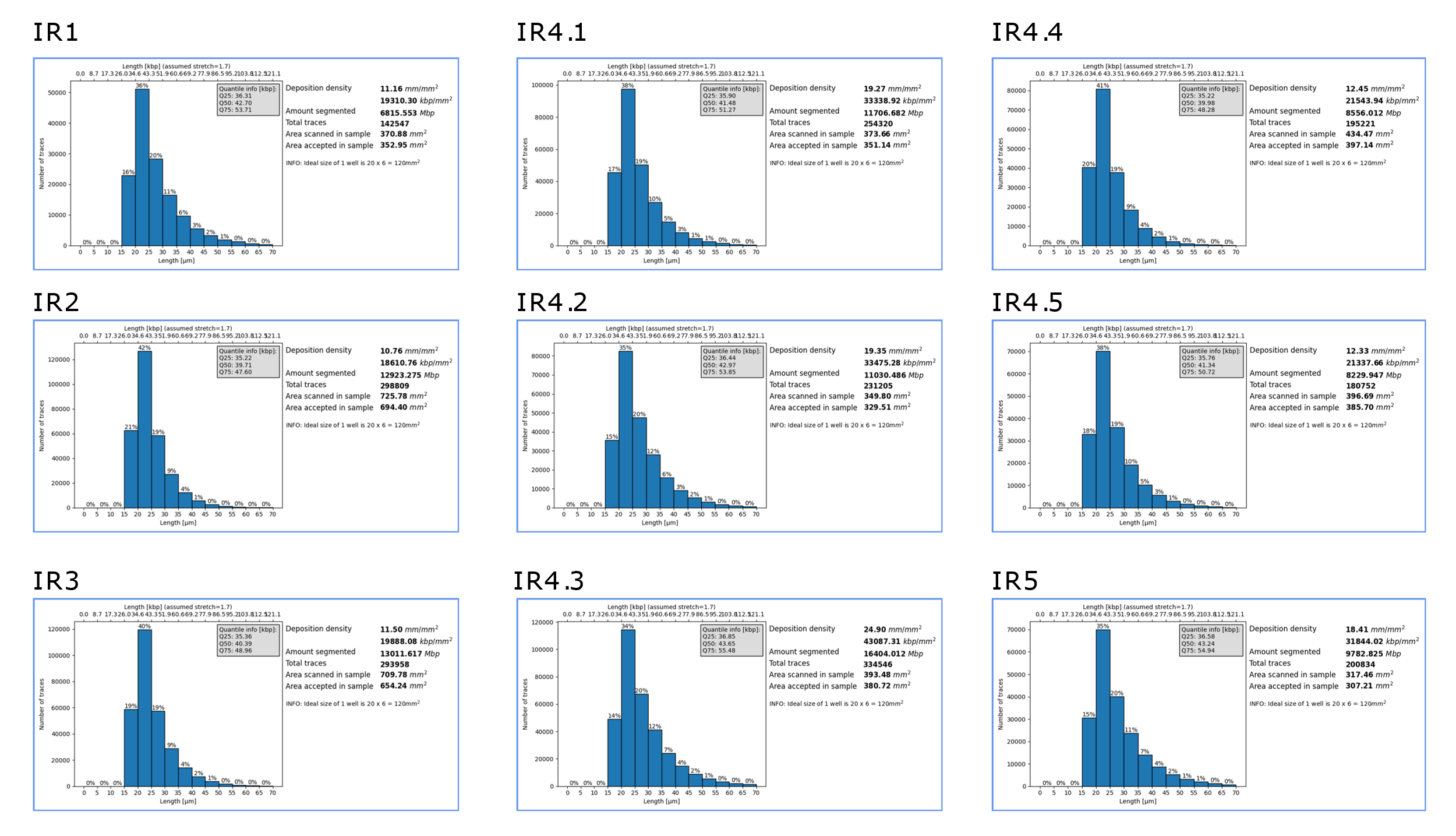
**Supplementary Figures**

Figure S2: The representative distribution of stretch factors from fecal dataset IR5, showing a peak at 1.75-1.76, and a heavy tail going as low as 1.64.

Figure S1: Sample summaries of all fecal optical mapping datasets, including the length histograms (assuming a stretch factor of 1.7). The summary statistics. Q25, Q50, Q75 and the number of traces is shown in the insets.

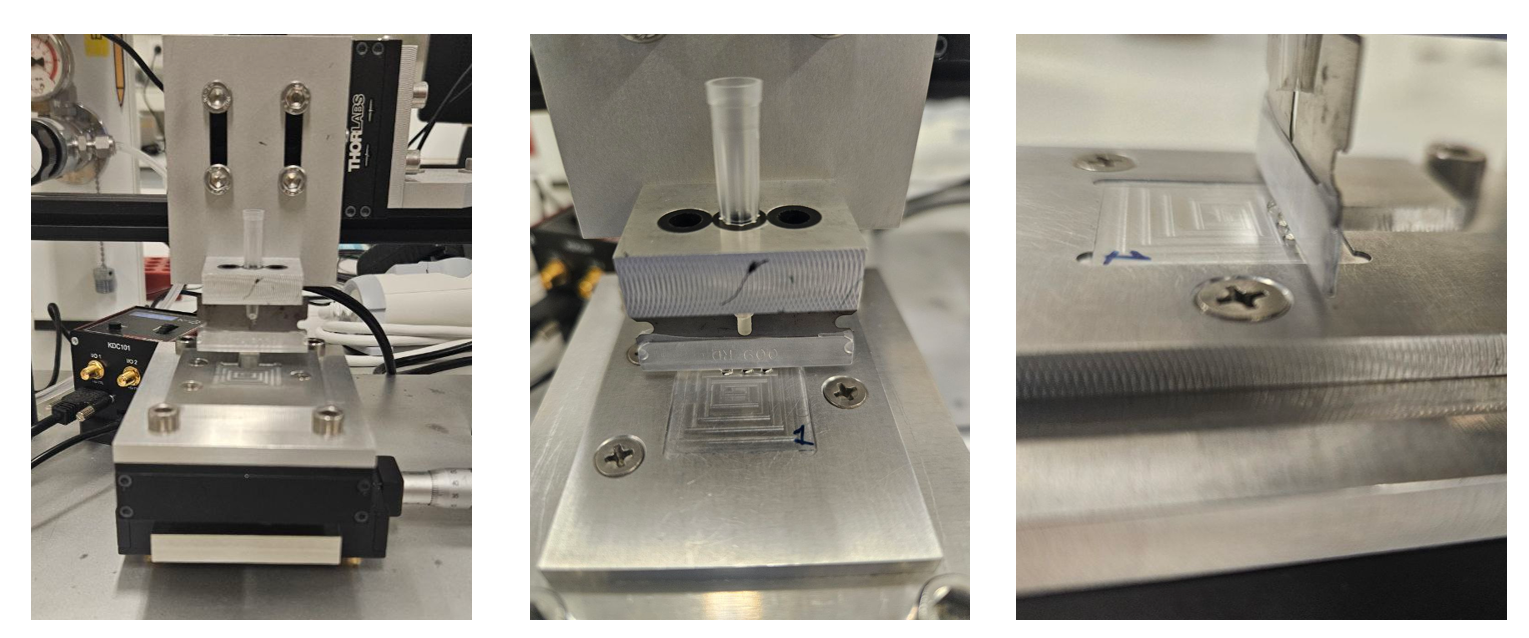

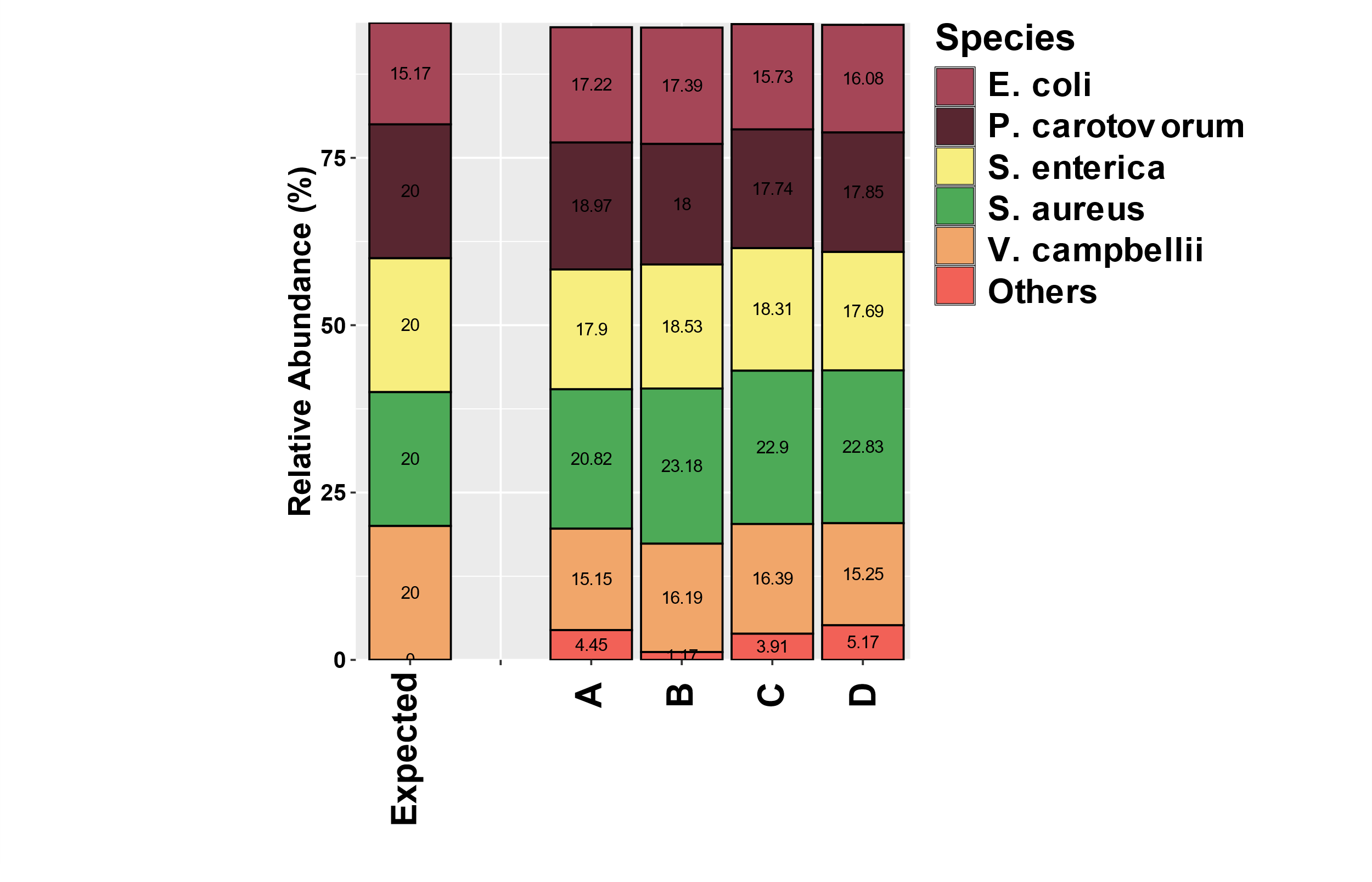

Figure S4: Images of the combing setup. The micrometer is adjusted to heigh of 0.85 mm above the coverslip and mounted on a translation stage. The stage holding the coverslip is driven by KDC101 motor, with a speed of 0.7 mm/s

Figure S3: Abundances of 5 species mixture from main text Figure 1b, including the false positives (denoted as ‘Others’)

**Supplementary Note 1: Error propagation on ALR transformed abundances**

To study the influence of optical mapping solely, in absence of any biases, we’ve created our own mock community by combining HMW DNA from 5 bacterial cultures together, with equal abundances for all, based on their Qubit measured concentrations. There are 2 possible sources of experimental error that may skew the abundance estimation using optical mapping: the pipetting error and the error stemming from concentration measurements. To take these into account, we’ve decided to estimate the possible theoretical errors on the absolute log-ratio abundances of our mixture, assuming effects stemming only from volume or concentration measurements. Consider the ALR of the present genomic content of genome $g_{i}$ with respect to $g_{n}$, defined as:

$$ALR\left( g_{i} \right)=\ln\left( \frac{g_{i}}{g_{n}} \right)=\ln\left( \frac{c_{i}v_{i}}{g_{n}} \right)$$

The error on ALR can be propagated as:

$$\sigma_{ALR\left( g_{i} \right)}= \sqrt{\left( \frac{\partial ALR}{\partial c_{i}}\sigma_{c_{i}} \right)^{2}+\left( \frac{\partial ALR}{\partial V_{i}}\sigma_{V_{i}} \right)^{2}{+\left( \frac{\partial ALR}{\partial g_{n}}\sigma_{g_{n}} \right)}^{2}}$$

Since $g_{i}=c_{i}V_{i}$, we can calculate the partial derivatives as:

$$\frac{\partial ALR}{\partial c_{i}}=\frac{\partial}{\partial c_{i}}\ln(\frac{c_{i}V_{i}}{g_{n}})=\frac{1}{c_{i}} ;\frac{\partial ALR}{\partial V_{i}}=\frac{\partial}{\partial V_{i}}\ln\left( \frac{c_{i}V_{i}}{g_{n}} \right)=\frac{1}{V_{i}}$$

Substituting, we obtain:

$$\sigma_{ALR\left( g_{i} \right)}=\sqrt{\left( \frac{\sigma_{c_{i}}}{c_{i}} \right)^{2}+\left( \frac{\sigma_{V_{i}}}{\partial V_{i}} \right)^{2}{+\left( \frac{\partial ALR}{\partial g_{n}}\sigma_{g_{n}} \right)}^{2}}$$

The last derivative can be obtained as:

$$\frac{\partial ALR\left( g_{n} \right)}{\partial g_{n}}= -\frac{1}{g_{n}}$$

And the standard deviation of $g_{n}$:

$$\sigma_{g_{n}}=\sqrt{\left( \frac{\partial g_{n}}{\partial V_{n}}\sigma_{V_{n}} \right)^{2}+\left( \frac{\partial g_{n}}{\partial c_{n}}\sigma_{c_{n}} \right)^{2}}=\sqrt{\left( c_{n}\sigma_{V_{n}} \right)^{2}+\left( V_{n}\sigma_{c_{n}} \right)^{2}}$$

Therefore, the final error on ALR yields:

$$\sigma_{ALR\left( g_{i} \right)}=\sqrt{\left( \frac{\sigma_{c_{i}}}{c_{i}} \right)^{2}+\left( \frac{\sigma_{V_{i}}}{\partial V_{i}} \right)^{2}{+\left( \frac{\left( c_{n}\sigma_{V_{n}} \right)^{2}+\left( V_{n}\sigma_{c_{n}} \right)^{2}}{g_{n}^{2}} \right)}}$$

The errors on the concentrations were determined empirically by 5 repeated measurements on the Qubit machine, and are given in Supplementary Table 1:

Supplementary Table 1: The concentration (ng/µl) measurements for each of the DNA stocks constituting the 5-bacteria mock mixture used for determination of $\sigma_{c_{i}}$

| Species | Measurement 1 | Measurement 2 | Measurement 3 | Measurement 4 | Measurement 5 |
| --- | --- | --- | --- | --- | --- |
| *E. coli* | 20,7 | 21,3 | 21,6 | 23,3 | 23 |
| *S. enterica* | 18,8 | 20,4 | 16,7 | 20,6 | 20,1 |
| *S. aureus* | 15,6 | 15,8 | 24,6 | 15,8 | 15,4 |
| *P. carotovorum* | 14,9 | 14,9 | 14,6 | 14,2 | 14,5 |
| *V. harveyii* | 5,64 | 5,48 | 5,88 | 5,8 | 5,9 |

The error on the volume was obtained from the pipet calibration, and yielded $V_{err}=0.099 \mu l$ for *E. coli, S. enterica, S. aureus, P. carotovorum* and $V_{err}=0.050 \mu l$ for *V.harveyii* respectively*.*

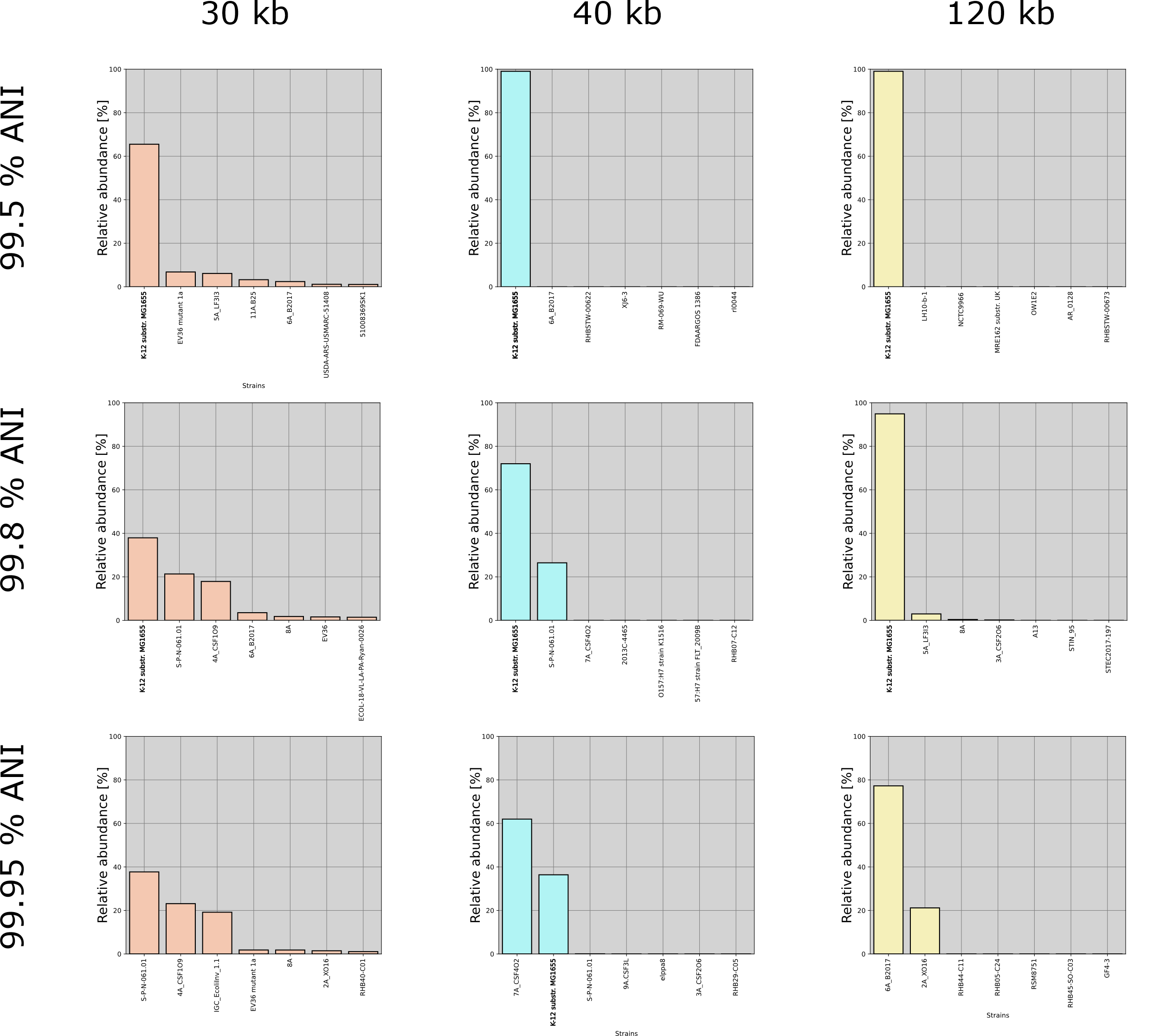
**Supplementary Note 2: Strain abundance calculation by maximum likelihood estimation from simulated *E. Coli str. K12 MG1655* data.**

Figure S5: Abundances of strains for datasets of 3 various lengths, matched against databases containing strains up to 3 different ANI thresholds with respect to the ground truth (GT) strain. The GT strain is indicated in bold (K12 substr. MG1655).

To assess the limit of resolution of optical mapping in presence of multiple strains, we’ve performed a controlled experiment on simulated data from *E. Coli str. K12 MG1655.* We expected that the proper strain abundance estimation would depend primarily on 2 factors: divergence between strains in the database, and the length of available optical maps. We therefore simulated 1000 maps, for 4 datasets of lengths of 30,40 and 120 kb, with a stretch factor of 1.75 and labelling efficiency of 75%. Next, we prepared 3 databases of *E. coli* strains, which contained *E. coli* genomes of complete a chromosome level assembly from NCBI, that are up to 99.5%, 99.8% and 99.95% similar to *E. coli MG1655* respectively. The final resulting databases contained 2914, 4102 and 4144 strains of *E. coli.* We then applied our alignment pipeline for each dataset with 3 databases and calculated the strain level abundances using our MLE approach.

In most cases, the GT strain is recovered with the highest abundance, however, in case of the 30 kb dataset a significant amount of cross assignment is observed. This could be potentially due to the lack of contextual information at that length. At 40 kb, the GT strain could be recovered from a database containing strains of up to 99.8% ANI, however a single cross match is observed to strain S-P-N-061.01 (NCBI accession number CP092698.1), which shares 99.67% ANI with the GT strain. In case of the highest length however, even at 99.8% ANI it was possible to correctly find the GT strain. At 99.95% ANI, none of the lengths were sufficient to recover the abundances of the GT strain, indicating a limit of taxonomic detection for optical mapping.

The database of D6300 strains consisted out of strains with an assembly level of ‘complete’ or ‘chromosome’ for strains obtained from NCBI. The total number of strains for each species is available in Supplementary Table 2:

| Species | Number of strains |
| --- | --- |
| *Escherichia coli* | 4428 |
| *Salmonella enterica* | 2143 |
| *Staphylococcus aureus* | 1634 |
| *Enterococcus faecalis* | 563 |
| *Bacillus spezizenii* | 7 |
| *Pseudomonas aeruginosa* | 794 |
| *Listeria monocytogenes* | 330 |
| *Lactobacillus fermentum* | 57 |

Supplementary Table 2: Total number of strains per species of ZymoBIOMICS D6300 used for taxonomic resolution experiment.

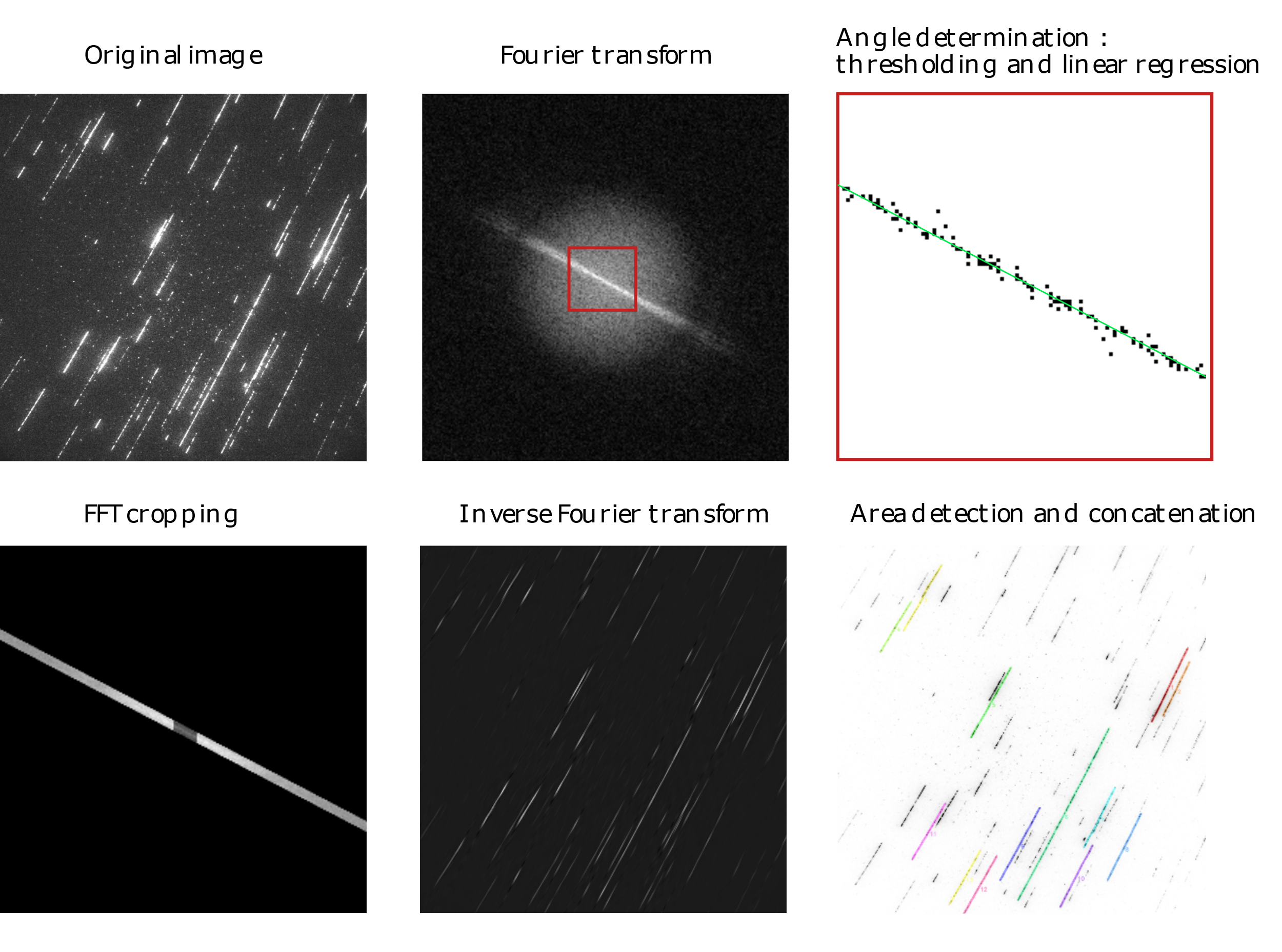
**Supplementary Note 3: Image processing pipeline**

Figure S6: An illustration of area joining rules in order to obtain a proper concatenation of traces. The total area available for joining is denoted in yellow.

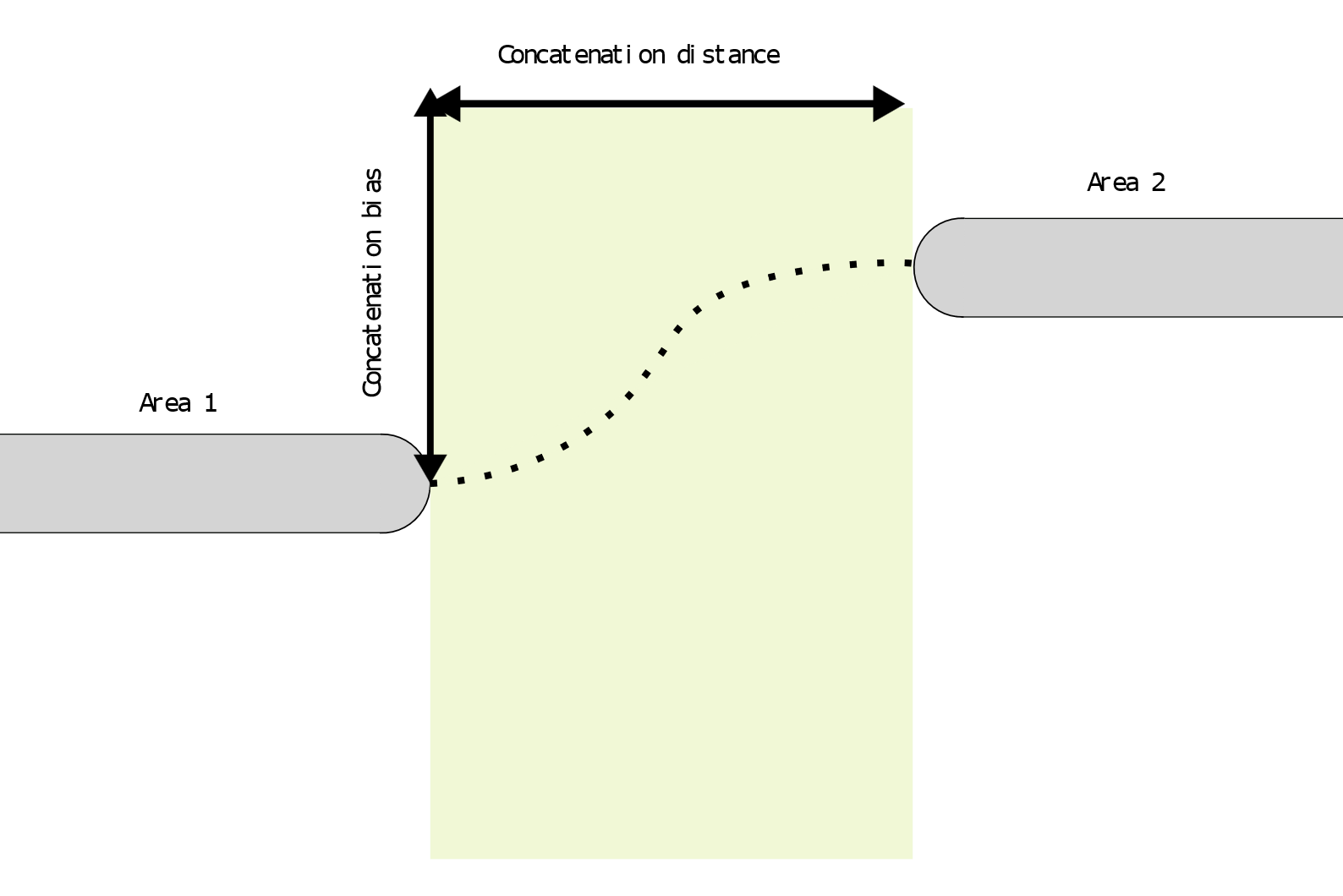
The obtained areas after thresholding and labelling the original image needed to be concatenated, as to obtain longest possible optical maps, with a low degree of interruption. These potential breakpoints occur due to inhomogeneous distribution of labels within the optical maps, or a non-linear stretching/curvature of the optical map on the Zeonex slide. To join the detected areas, we’ve imposed an empirical criterion on the location of endpoints of both optical maps as follows: two optical maps are joined together only if their ends are shifted within a concatenation distance of one another (7 pixels) in the x-direction and displaced no more than a concatenation bias of 8 pixels far in the y-direction.

Figure S7: The image processing pipeline, with a visualization of every step, starting from the original image and ending with the final labelling of individual traces using the obtained mask.

**Supplementary Note 4: Multi-scale cross-correlation**

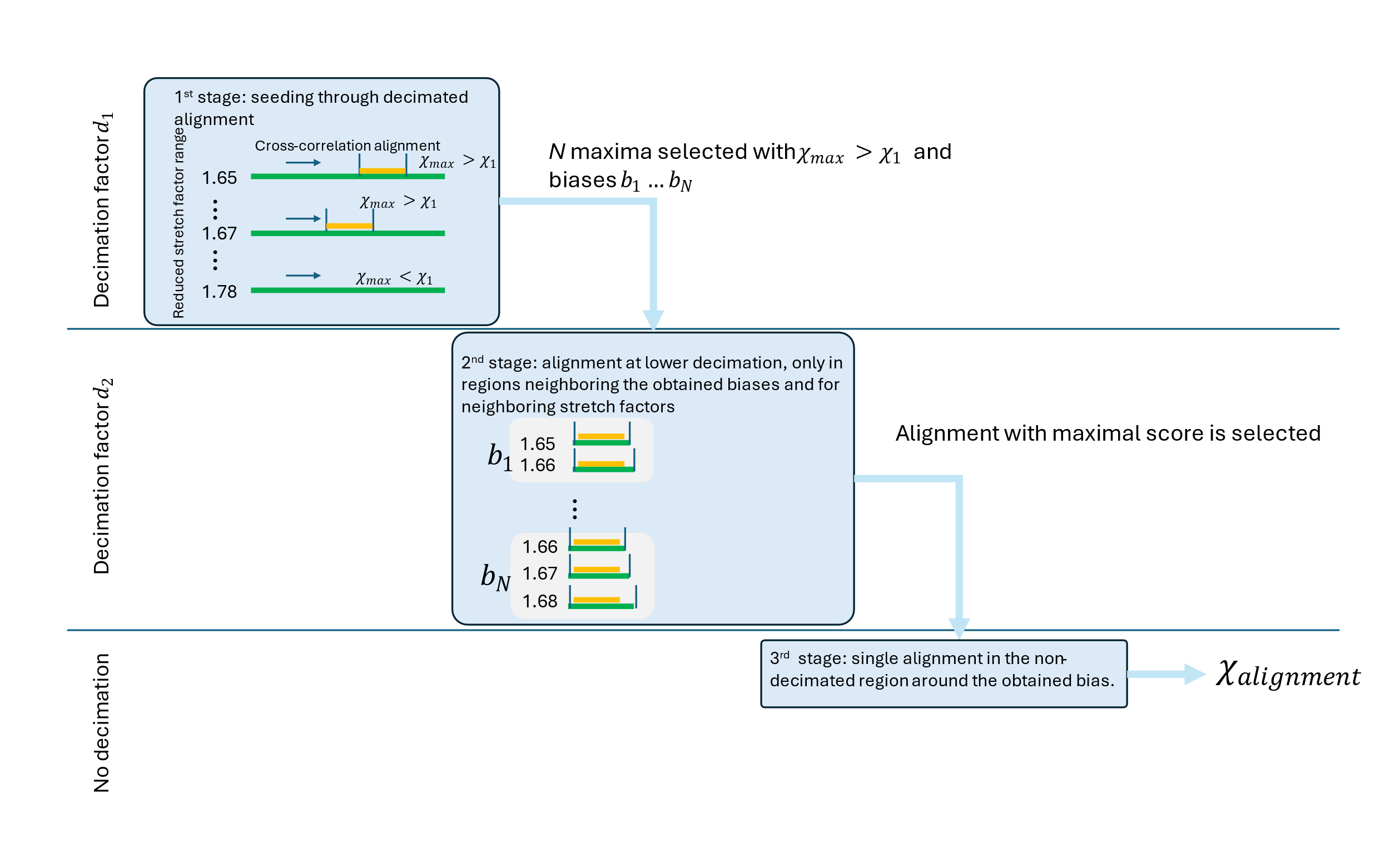
Assignment of optical maps to the reference database becomes challenging with growth of the database. Therefore, we attempted to speed up the classification performance through modification of the cross-correlation algorithm. Starting from a query of length *L* and a reference of length *N,* the time complexity of cross-correlation in time domain is given as $O\left( NL \right)$ and in frequency domain as $O\left( N\log N \right)$, when using fast Fourier transform for cross-correlation computation. However, since we need to perform the cross-correlation for multiple stretch factors, the complexity needs to be multiplied by the number of stretch factors $s$, yielding $O\left( sN\log N \right)$.

Figure S8: The outline of the multi-scale cross-correlation algorithm, showing the 3 step process.

In the first step of our optimization, we’ve down sampled both the query and the reference by the factor $d_{1}=4$ . Furthermore, we’ve decimated the number of stretch factors by taking every second stretch factor. With these adjustments, the first step yields a reduction the time complexity, with the time complexity being $O\left( \frac{s}{2}\frac{N}{d_{1}} \log\frac{N}{d_{1}} \right)$. For each alignment, the bias of all cross-correlation scores exceeding a threshold value of $\chi_{1}=0.5$ is recorded and remaining *N=3* maxima with biases $b_{1}\ldots b_{N}$ and stretch factors $s_{1}\ldots s_{N}$are selected to be processed in the next step. If no maxima exceeding the threshold of 0.5 are found, the procedure is aborted.

In the second step, the cross correlation is computed on data which has been down sampled by a factor of $d_{2}=2$ and only between the down sampled query and the down sampled reference in regions around the biases obtained from the previous step. In this case, a portion of the down sampled reference trace, ranging from $b_{i}\frac{d_{1}}{d_{2}}-\delta_{1}$ to $b_{i}\frac{d_{1}}{d_{2}}+L/d_{2}+ \delta_{1}$ is taken, where $\delta_{1}=10$ is the offset from the obtained bias $b_{i}$ and a cross-correlation is performed only within these boundaries. Additionally, a cross-correlation alignment is also performed for the neighbouring stretch factors $s_{i}-\delta s$ and $s_{i}+\delta s$, with $\delta s=0.01$. From all the alignments, a single alignment corresponding to the maximum cross-correlation value from all alignments is selected for the next step, together with its bias $b_{max}$and stretch factor $s_{max}$values.

In the last step, the cross-correlation is performed between non decimated data in the region corresponding to the $b_{max}d_{1}-\delta_{2}$ and $b_{max}d_{1}+\delta_{2}$ of the original reference, where $\delta_{2}=20$ is the offset defined for this step. The final alignment yields the final alignment score $\chi_{alignment}.$Notably, since for the last 2 steps, the alignment is only performed within the seeded region originating from step 1, the computational complexity of these steps is practically negligible in comparison to the first step.

Although the maximum cross-correlation score can be used as a final threshold, this is suboptimal [1]. Therefore, we introduced an additional scoring metric, which compares the cross-correlation score to the distribution of possible scores that are sampled empirically. In order to get the underlying distribution, we have generated a single random fragment, with length of 5 Mb. Next, we perform a single cross-correlation alignment and sample all cross-correlation scores from all lag positions. This yields a distribution, which can be approximated by a Gaussian with a zero mean, which allows us to calculate the standard deviation $\sigma_{\chi}$ for each experimental optical map. To gauge the quality of alignment, we introduce the $q$- score, which is defined as: $2^{-\frac{1}{10}\frac{\chi_{max}}{\sigma_{\chi}}}$, which is essentially a transformed Z-score assuming a zero mean. The optical map alignments are thresholded at $q_{ths}=0.635$ for all datasets. The threshold represents a compromise between TP and FP rate, which was determined from performance on D6300 mock data (see Figure 1c), achieving an FP rate between 1-4% relative abundance.
